# Boosting the speed and accuracy of protein quantification algorithms in mass spectrometry-based proteomics

**DOI:** 10.1101/2025.10.06.680769

**Authors:** Thang V Pham, Chau TM Tran, Alex A Henneman, Long HC Pham, Duc G Le, An H Can, Phuc HL Bui, Sander R Piersma, Connie R Jimenez

**Affiliations:** Amsterdam UMC, location Vrije Universiteit Amsterdam, Department of Medical Oncology, OncoProteomics Laboratory, Amsterdam, the Netherlands; Cancer Center Amsterdam, Imaging and Biomarkers, Amsterdam, the Netherlands; University of Engineering and Technology, Vietnam National University, Hanoi, Vietnam

## Abstract

Current methods for protein level quantification in mass spectrometry-based proteomics do not scale with the increasing number of samples because of limited system memory and algorithmic complexities. Here we propose a new data structure that supports parsing of input as data stream, improve state of the art quantitation methods in performance by orders of magnitudes, and provide a generic method to boost the precision and accuracy by fragment weighting.

## Main

Protein quantification or summarization is a crucial processing step in mass spectrometry-based proteomics that combines quantitative values at peptide fragment ion level into protein level quantification. With the increase in the number of samples driven by advances in automated sample preparation, for example a study with 36,000 samples [1], the current data processing methods become inefficient or fail to complete due to limitations in system memory and inherent algorithmic complexity. The state of the art MaxLFQ algorithm [6], as implemented in the R package *iq* [13], becomes computationally inefficient for large datasets, requiring over a day to process 10,000 samples and failing to complete for 25,000 samples on a standard workstation, which highlights the need for new method development [2]. Here we present new implementations of protein quantification methods, including MaxLFQ, that are several orders of magnitude faster than current algorithms, enabling the processing of hundreds of thousands of samples.

To handle a very large dataset, we developed a new data structure that avoids the need to load all data into memory for computation. The input for protein quantification, which is a data table generated by raw data processing tools like DIA-NN [7], becomes too large to be held in memory as the number of samples increases. Fig. 1a describes the new data structure, which consists of three separate indexing files for protein, sample, and fragment ion identifiers. The quantitative values for each protein are stored in a dedicated data file that references these sample and fragment indices. This data structure is efficient as it eliminates duplications in protein, sample, and fragment description. More importantly, processing the input as a data stream alleviates the need for large memory, and this approach also facilitates distributed computing by allowing subsets of proteins to be processed on different nodes.

**Fig. 1.**
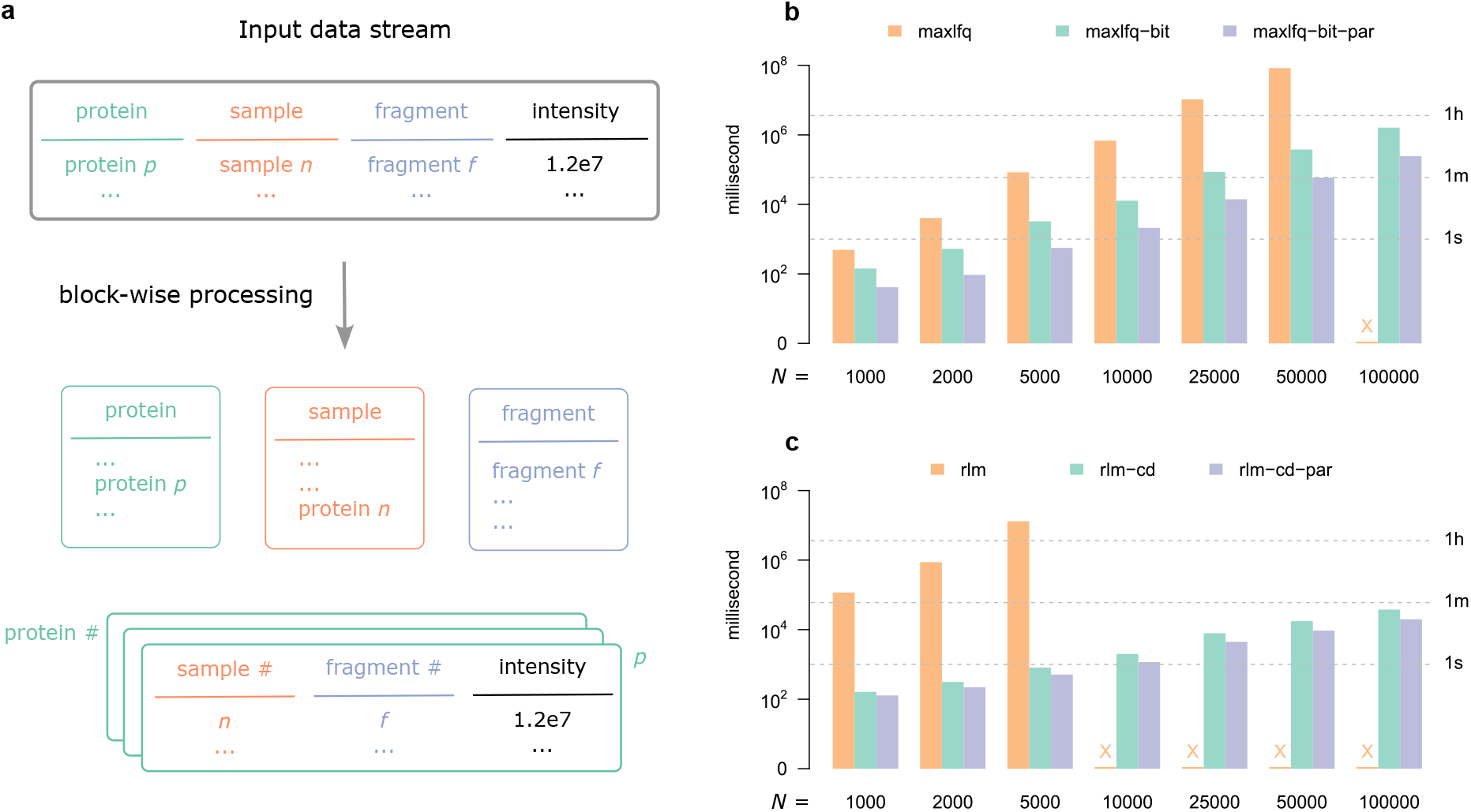
Optimizing data structure and algorithm for protein quantification. **a**, a new data structure that keeps proteins, samples, and fragment ions in index files and quantitative intensities in individual files. A data stream can be processed in a block-wise manner. **b**, comparing execution time of the current state of the art MaxLFQ algorithm (maxlfq) versus a bit matrix-based implementation (maxlfq-bit), and its parallelized version (maxlfq-bit-par) with respect to sample sizes on a simulated dataset. **c**, comparing execution time of the robust linear model (rlm) versus a coordinate descent implementation (rlm-cd) and its parallelized version (rlm-cd-par).

Next, we addressed the challenge of scaling up quantification algorithms, as memory is a limiting factor, even for a single protein due to algorithmic complexity. One of the most popular methods is the MaxLFQ algorithm that addresses missing values and differing ionization efficiencies by solving an optimization problem based on ratios between pairs of samples. As a result, it has quadratic complexity due to the consideration of all sample pairs, which makes it computationally inefficient for a large number of samples. Specifically, it requires solving a linear system of equations involving an (*n* + 1) × (*n* + 1) matrix where *n* is the number of samples (see Methods). Forming this matrix in memory is a limiting factor for large *n*. However, we have discovered that this matrix can be represented as the sum of a binary matrix and a diagonal matrix, which enables a highly efficient implementation. By combining a bitwise representation of the matrix and the conjugate gradient descent algorithm [10], we can quantify a simulated example of 1 million samples in under 8 hours. As shown in Fig. 1b, at *n* = 50,000, our new implementation (maxlfq-bit) is more than two orders of magnitude faster than the current MaxLFQ implementation (maxlfq), while its parallel version (maxlfq-bit-par) is more than three orders of magnitude faster on an 8-core CPU. The current MaxLFQ implementation failed at *n* = 100,000. In contrast, the maxlfq-bit version finished in under 30 minutes, and the parallel maxlfq-bit-par version completed in just 4 minutes.

We examined three other methods published in the literature, the median polish method [16] used by the popular R package *MSstats* [5], the msqrobsum method used by the package *msqrob2* [15], and the directLFQ algorithm [2] designed for scalability. The median polish and directLFQ are two computationally efficient methods. The msqrobsum method which optimizes a robust linear model fails at *n* = 10,000 (Fig. 1c). A close investigation shows that the method solves a series of least square problems, where the matrix involved is a sparse matrix, and hence a method like conjugate gradient can be used. However, motivated by the efficiency of the coordinate descent approach [9], we implemented an algorithm that minimizes the objective function one variable at a time. This algorithm is highly efficient and easy to parallelize, processing one million samples in just over 4 minutes. As shown in Fig. 1c, the new single-threaded implementation completes the analysis for *n* = 100,000 in under one minute. Hence, all three methods are faster than MaxLFQ.

We conducted a brief evaluation of the quantification accuracies of the different methods to assess whether the computational effort of MaxLFQ is justified. In addition, we investigated if incorporating fragment weighting will improve quantification accuracy by reducing the influence of noisy fragments. To this end, we employed a generic weighting approach, as illustrated in Fig. 2a. We computed an initial estimate and subsequently employed a distance metric that accounted for missing values to measure the difference between each fragment and the initial estimate [8]. Fragments that were in proximity of the initial estimate had weights of 1 while others had weights inversely proportional to their distances. The weights derived from the Huber method [11] were then used in a second round of quantification (see Methods).

**Fig. 2.**
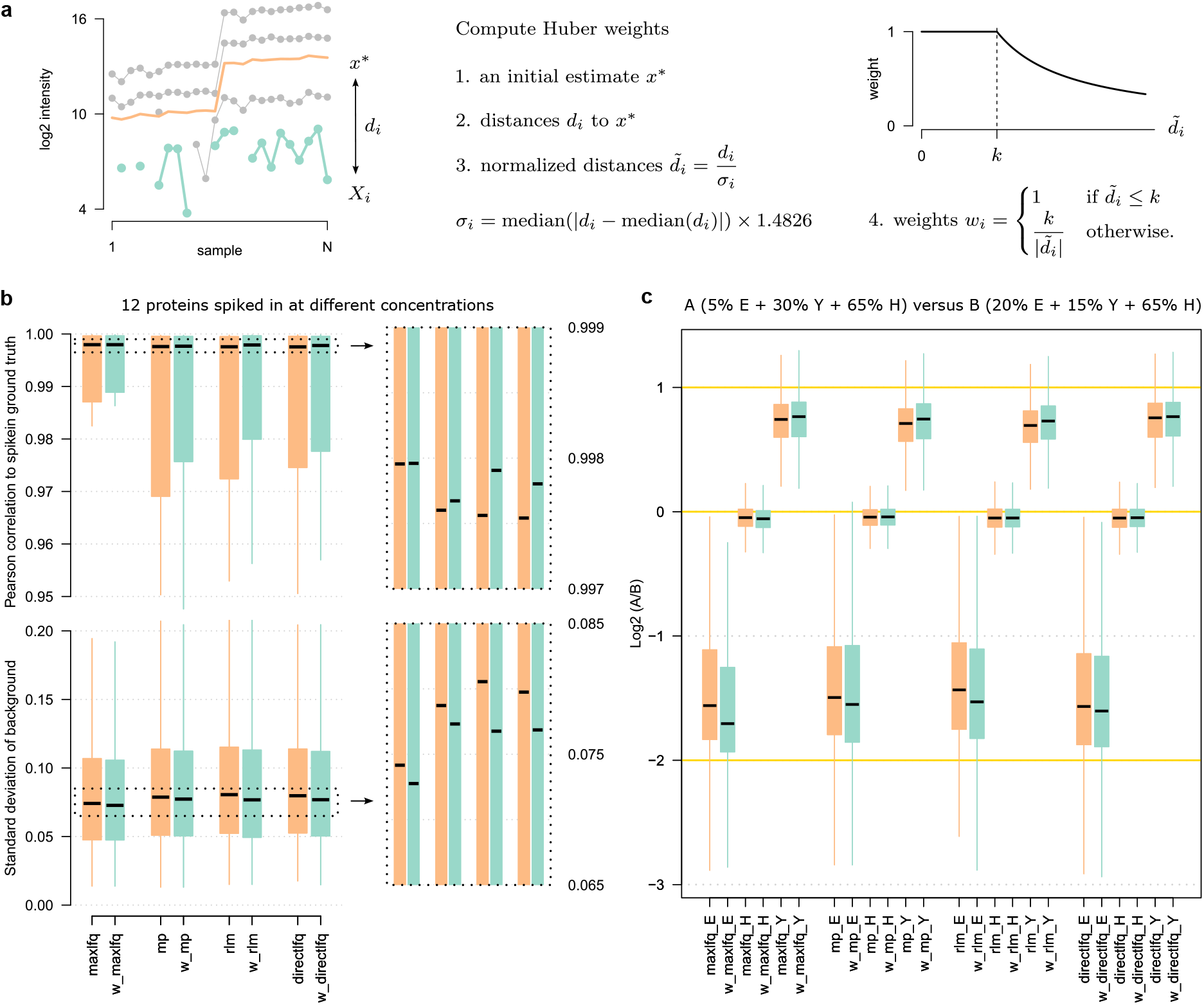
Weighting fragment ions improves quantification accuracy. **a**, the Huber weighting method consisting of four algorithmic steps. The left sub-panel shows raw data in gray with a fragment highlighted in light cyan, and an initial estimate in light orange. **b**, boxplots of the performance of four methods, MaxLFQ (maxlfq), median polish (mp), robust linear model (rlm), directLFQ (directlfq), and their weighted version (with the prefix w) on a dataset with 12 proteins spiked in a stable background at different concentration. Upper panel, correlation of the spike-in to ground truth values (higher is better). Bottom panel, variation of the background proteins (lower is better). **c**, performance on the mixed species dataset where yeast and *E. coli* proteins are spiked in a human protein background at different ratios. The golden lines are the expected log2 ratios, -2 for *E. coli* proteins, 0 for human proteins, and 1 for yeast proteins. The results for the three species have postfix _E, _H, and _Y for *E. coli*, human, and yeast respectively.

We evaluated the base methods and the reweighting strategy on two datasets. Fig. 2b shows the result of the four algorithms and their weighted versions on a dataset with 12 proteins spiked in at different concentrations to a common background [4]. Here all weighted versions outperform the base methods in both correlation to the theoretical spike-in concentration and the stability of background proteins. Among the base, non-weighted methods, the MaxLFQ algorithm outperform others, justifying its popularity despite of the computational effort. The weighted versions consistently outperform the non-weighted versions. Fig. 2c shows the result on a mixed species dataset [17] in which *E. coli* and yeast are mixed with a human protein background at two different concentrations, create three theoretical ratios for comparative analysis, equal for human background, 4-fold for *E. coli* and 2-fold for yeast. Again, the MaxLFQ algorithm out-performs other base methods while the weighted methods outperform the base methods in all cases.

In conclusion, the maxlfq-bit and rlm-cg implementations and their parallelized versions are the method of choice for the MaxLFQ algorithm and the robust linear model, respectively. These methods are available in the R package *iq* version 2. The source code is publicly available under a permissive license, allowing for straightforward integration into existing software solutions. Furthermore, the use of fragment ion re-weighting is highly recommended, as it substantially improves the accuracy of all algorithms.

## Methods

### A fast MaxLFQ algorithm

Let *X* be an *m* × *n* data matrix of log2-transformed intensities for a single protein, where the *m* rows represent quantified fragments or peptides, and the *n* columns represent samples. The matrix *X* may contain missing values. The protein quantification problem is to estimate an *n*-vector ***x*** for relative protein abundance across all *n* samples. The MaxLFQ approach obtains ***x*** by minimizing the sum of squared differences between the estimated pairwise sample ratio and the median of the observed ratios [6]. The algorithm in [13] solves this optimization problem by forming an equality constrained least squares, and subsequently ***x*** is obtained as a solution of a system of (*n* + 1) linear equations

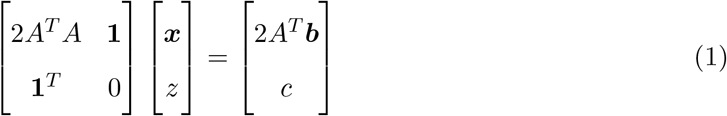

where *A* is a *p* × *n* matrix, ***b*** a *p*-element vector containing all *p* pair-wise sample ratios, **1** an *n*-vector of ones, *z* an auxiliary variable, and *c* a scaling constant. Each row of *A* contains all zeroes except for two entries with value of −1 and +1 corresponding to the sample ratio in ***b***. The linear system of equation was solved using a standard library [3]. The (*n* + 1) × (*n* + 1) matrix on the left-hand side needs to be computed and stored in memory, which limits the analysis to about 10,000 samples for a workstation with 128 GB of RAM.

In a new method, we employ the conjugate gradient algorithm [10] to solve (1), which requires repeated multiplication of the fixed (*n* + 1) × (*n* + 1) matrix and different (*n* + 1)-vectors. Essentially, the matrix does not need to be stored in memory; it can be constructed on demand. However, reconstructing the matrix is computationally expensive. The key idea is that this matrix can be decomposed into a binary matrix and a diagonal matrix, leading to a memory-efficient bit-based representation. This enables keeping a square matrix exceeding 1 million by 1 million in memory on a workstation with 128 GB of RAM. This method is referred to as maxlfq-bit. A parallel implementation is referred to as maxlfq-bit-par.

### A fast robust linear model algorithm

Under an ideal, error-free condition, each fragment intensity is proportional to the protein intensity. That is, each row of *X* is equal to ***x*** plus a scaling term in the log2 space. Let ***s*** be the vector of *m* scaling terms, one for each fragment. The matrix *X* is then the sum of two matrices. The first is formed by replicating ***x*** row by row, and the second replicating ***s*** column by column. We can estimate ***x*** and ***s*** by minimizing the difference between the observed and theoretical values

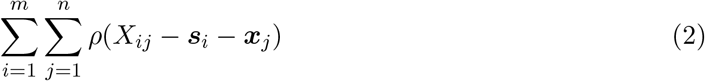

where *ρ* is a penalty function. The msqrobsum method [15] uses the Huber loss as its penalty function [11]. The method is implemented in the function robustSummary in the R package *MsCoreUtils* [14], calling the rlm function in the R package *MASS* [18]. The rlm function forms a matrix of size (*m* + *n*) × (*m* + *n*) and, hence, is not efficient for large sample size due to memory constraints. We implemented a new algorithm that uses coordinate descent to optimize one variable at a time. Similar to the elastic net algorithm [9], our approach to solving (2) by minimizing individual coordinates is both computationally and memory-efficient. Since we do not use the experiment design feature of msqrobsum, we refer to the base algorithm as robust linear model (rlm), the coordinate descent implementation as rlm-cd, and its parallelized version as rlm-cd-par.

### The median polish algorithm

The median polish algorithm [16] is a very fast method to obtain ***x*** and ***s*** by alternately subtracting the row and column medians of the residual matrix in equation (2). It can be shown that the median polish algorithm decreases the value of the objective function in (2) in each iteration when *ρ* is the absolute function. The algorithm is available in the R package *stats* which is single-threaded. We reimplemented the algorithm in C++ to take advantage of parallel processing. We set the convergence criteria to a tolerance of 1e-4 and a maximum number of iterations of 1000.

### The directLFQ method

The directLFQ algorithm was designed for scalability to a large number of samples [2]. We used the directLFQ version 0.3.2. We converted the input data to the directLFQ format and ran the software with the minimum number of ion intensities necessary set to 0 (`--min_nonan0`) and no normalization (`--deactivate_normalization True`) as the normalization was done prior to the format conversion.ss

### Adapting quantification methods for fragment weights

For the MaxLFQ algorithm, we used weighted median in the calculation of pair-wise sample ratios ***b*** in (1). For the median polish algorithm and the robust linear model, we incorporated the weight factors in the optimization criterion (2). For the directLFQ method, we resampled with replacement the fragments 100 times according to the weights to create 100 input datasets, ran the algorithm 100 times, and subsequently averaged the results.

### Weighting fragments

We ran each base algorithm twice. The first run led to an initial estimate ***x***^∗^. For each fragment (row *i* of *X*) we calculated coordinate-wise distances to ***x***^∗^, taking into account missing value as follows

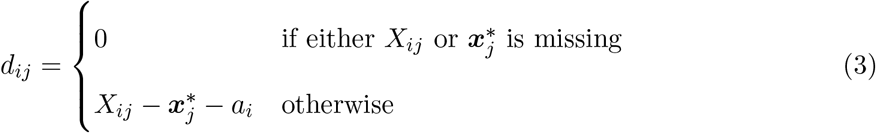

where *a*_*i*_ is an alignment term to align fragment *i* to ***x***^∗^. Subsequently, the distance between fragment *i* and ***x***^∗^ is

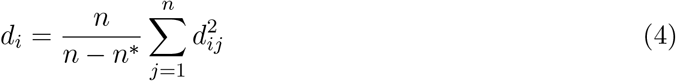

where *n*^∗^ denotes the number of missing values. Next, we computed a standardized differences by dividing the distances to the median of absolute deviation times a scaling factor

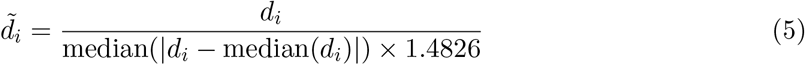

This derivation was motivated by the Huber weighting [11]. Finally, the weights were given by the Huber weight

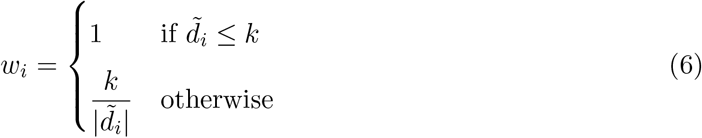

with the constant *k* = 1.345. Intuitively, fragments that are further from ***x***^∗^ will get smaller weights, whereas others will have an equal weight of 1.

### Datasets

#### The Bruderer15 dataset

The dataset in [4], reprocessed by Spectronaut version 13.0, was downloaded from the *iq* package GitHub repository (see Data availability). The dataset contains mass spectrometry analysis of 8 biological samples in which 12 proteins were spiked in at different concentrations. There are 3 technical replicates for each biological sample. Thus, there are 24 samples in total.

#### The VanPuyvelde22 dataset

We processed Bruker timsTOF Pro data from [17], downloaded from the ProteomeXchange repository PXD028735. This is a mixed species dataset of 27 samples in which condition A composed of 5% *E. coli*, 30% yeast, and 65% human and condition B 20% *E. coli*, 15% yeast, and 65% human. We used DIA-NN [7] version 2.2.0 with precursor and fragment m/z ranges of m/z 399-1201 and m/z 50-2000 setting as described in [12]. Both mass accuracy and MS1 accuracy were set to 15ppm. The options `--export-quant` and `--no-peptidforms` were switched on. Ox(M) was selected and maximum number of variable modifications was set to 1. For database searches, we used canonical Swiss-Prot subsets of the UniProt reference proteomes as of 16.05.2025 (human - UP000005640 with 20647 entries; yeast - UP000002311 with 6065 entries; *E. coli* - UP000000625 with 4402 entries), and the MaxQuant contaminant database. The raw intensities were normalized by the median of the human proteins.

### Execution time analysis

All timing experiments were carried out on a workstation with 128 GB RAM, 8 cores Intel Xeon w3-2525 CPU running Microsoft Windows 11 and R version 4.5.1. We simulated a dataset with 1 million samples as follows. From the data matrix for one protein in the VanPuyvelde22 dataset (*E. coli* P0AEG4), we randomly sampled from the 27 columns, equally divided in three classes, and adding Gaussian noise (mean = 0, standard deviation = 0.001). The dimension of the simulated matrix is 56 rows × 1,000,000 columns. We took the first *n* columns from this simulated data matrix as required for the timing benchmark.

## Data availability

The Bruderer15 dataset can be downloaded from GitHub (https://github.com/tvpham/iq/ releases, data file Bruderer15-DIA-longformat-compact.txt.gz). The VanPuyvelde22 dataset and the simulated data can be downloaded from Zenodo (https://doi.org/10.5281/zenodo. 16929646).

## Code availability

The source code is available under the BSD-3-clause license on GitHub (https://github.com/tvpham/iq) and on the R package repository CRAN (https://cran.r-project.org/package=iq).

## Acknowledgements

This work has been supported by VNU University of Engineering and Technology under project number CN24.27 and by the Netherlands eScience Center project ASDI.2020.014.

